# Production Status, Breeding Priorities and Genetic Resources of Cowpea in Post-Civil War Somalia

**DOI:** 10.1101/2025.09.02.673644

**Authors:** Ahmed O. Warsame, Yahye A. Isse, Abdirizak Mohamed Sh. Abdi

**Affiliations:** Crop Science Centre, Department of Plant Sciences, University of Cambridge, Cambridge, UK; Faculty of Agriculture, Zamzam University of Science and Technology, Mogadishu, Somalia

**Keywords:** Cowpea, farmer survey, genetic resources, Somalia, food security

## Abstract

Over the past three decades, Somalia has faced recurrent civil conflicts and prolonged droughts, leading to widespread displacement of farming communities. These disruptions have weakened traditional seed systems and potentially damaged local crop diversity. In this context, to strengthen the country’s food security and climate change adaptation of smallholder farmers, there is an urgent need to revive agricultural research and breeding programs for the main food crops grown in the country. Cowpea (*Vigna unguiculata* (L.) Walp.) is the most important food crop after maize and sorghum in Somalia, providing inexpensive protein for humans and biologically fixed nitrogen for low-input crop production systems in the country. This study assessed the current status of cowpea production, identified breeding priorities across three agroecological zones in southern Somalia, and established a core germplasm collection for future breeding programs. Farmer surveys were conducted in Baidoa, Afgoie, and Jowhar, involving interviews with 150 farmers using a semi-structured questionnaire. The results showed that over 60% of farmers cultivate cowpea for both household consumption and market purposes. While the crop is predominantly grown as an intercrop, nearly 45% of farmers in Afgoie grow it as a sole crop. Marked regional differences were observed in varietal preferences, with farmers in Afgoie favouring erect, uniformly maturing cultivars with a dark-red seed colour, whereas other regions demonstrated more diverse preferences. Production constraints also varied by location, with drought identified as the primary limitation in Baidoa, and pests and diseases as the major challenges in Afgoie and Jowhar. Accordingly, farmers ranked early maturity as the top breeding priority in Baidoa, whereas disease and pest resistance were the primary targets in Afgoie. Furthermore, we established a germplasm collection representing Somalia’s cowpea diversity from both the pre- and post-war periods. The collection also includes a subset of global cowpea diversity and accessions with known desirable agronomic traits. These genetic resources, together with farmer survey data, provide a foundation for targeted breeding efforts aimed at improving cowpea productivity and enhancing food security in Somalia in the future.

## INTRODUCTION

As a result of the compounded effects of prolonged civil war and climate change on agricultural production, Somalia has frequently experienced food shortages, including two famine events in recent history (Majid & McDowell, 2012; Maxwell & Fitzpatrick, 2012; Moore et al., 1993). Currently, nearly a quarter of the population suffers from acute food shortages and malnutrition (Jamil et al., 2025). Therefore, an urgent step towards addressing the country’s chronic food crisis and vulnerability to climate change is to rebuild the crop production system by developing new crop varieties with improved climate resilience and desirable agronomic and quality attributes.

Cowpea (*Vigna unguiculata* (L.) Walp.) is the most important legume crop in Somalia and is mainly grown intercropped with maize or sorghum (Longley et al., 2001b; Padulosi & Fiore, 1989), making it the primary source of nitrogen in the low-input smallholder farming systems of Somalia. The crop is also cultivated as a sole crop in the central and southern regions along the coastal belt, where sandy soil and erratic rains make cowpea one of the few viable crops. In addition to its agronomic versatility, cowpea is a critical source of protein that enriches cereal-based diets in Somalia (Longley et al., 2001a; Padulosi & Fiore, 1989). For instance, the dinner dish called “Ambuulo”, a mixture of cowpea with imported rice or local cereals such as maize and sorghum, is consumed in nearly every rural and urban household. This local cowpea consumption generates an estimated annual gross value of $15.4 million (World Bank & FAO, 2018), making this crop crucial for the food security and economic development of smallholder farmers in Somalia.

The main challenge in reviving crop genetic improvement after 35 years of civil war is the lack of reliable data on the current status of crop production and the uncertainty regarding the condition of local crop genetic resources in the country. Before the civil war, the only documented official release of a cowpea cultivar was TVu-15, which was introduced to Somalia in collaboration with the International Institute of Tropical Agriculture (IITA) (Noor et al., 1983). However, it has been reported that this variety was less favourable to farmers because of its seed appearance (Rees et al., 1991). In late 1988 and early 1989, just a year before the collapse of the central government, a cowpea germplasm collection was conducted across Somalia (Padulosi & Fiore, 1989). A total of 92 accessions were collected and preserved in the IITA Genebank. Although the authors observed little diversity among local cowpeas based on their plant and seed morphology (size and colour), this collection is a valuable resource that represents Somalia’s historic cowpea genetic diversity. In a genetic diversity study of East African cowpea accessions, including 15 accessions from the Somali collection at IITA, Ayalew (2015) found that cowpea accessions from Somalia had a low polymorphic information content (PIC), and 13 of the 15 accessions formed a separate cluster. Considering the unique geographical and cultural nature of Somalia, this may indicate a long history of localised cowpea selection, resulting in a relatively narrow but distinct genetic base. It is unknown whether the local gene pool has shrunk further due to the prolonged civil war and climate change. In addition, international aid organisations frequently distributed emergency seed inputs to rehabilitate farmers displaced by conflict (Longley et al., 2001a, 2001b). Consequently, it is probable that some local cultivars were lost while new varieties were introduced, primarily from neighbouring countries, owing to the absence of domestic seed production capacity in Somalia (Longley et al., 2001b).

In this study, we assessed the production practices, constraints, and breeding priorities of cowpea in three regions of southern Somalia. We conducted a nationwide and worldwide cowpea germplasm collection to establish a core collection which captures the past and present state of Somalia’s cowpea diversity. This study lays the foundation for future breeding programs aimed at developing cowpea varieties with enhanced climate resilience and quality.

## MATERIALS AND METHODS

### Study sites

Farmer surveys were conducted across three regions in southern Somalia, namely Bay, Middle Shabelle, and Lower Shabelle (Figure 1). Within each region, a representative district was selected: Baidoa in Bay, Jowhar in Middle Shabelle, and Afgoie in Lower Shabelle. From each district, five villages were chosen, resulting in a total of 15 villages across all regions. These regions were selected to capture the diversity of agroecological conditions, livelihood strategies, and farming systems in southern Somalia, while considering accessibility and safety. The Bay region, often referred to as Somalia’s “sorghum basket”, is characterised by widespread sorghum cultivation, typically intercropped with cowpea (Longley et al., 2001a; Rees et al., 1991). The soils are predominantly clay, and agricultural production is entirely rainfed. Crop growth periods are relatively short, less than 90 days during the Gu’ season (April–June), and under 60 days during the Deyr season (October– December) (Venema, 2007). In contrast, some parts of the Middle and Lower Shabelle regions experience relatively longer growing seasons, with areas along the Shabelle River supporting irrigated commercial crop production, including the cultivation of maize and vegetables. The Jowhar district is part of the “cowpea belt”, which extends along the coastal regions of the Middle Shabelle, Galgudud, and Mudug in central Somalia.

**Figure 1.**
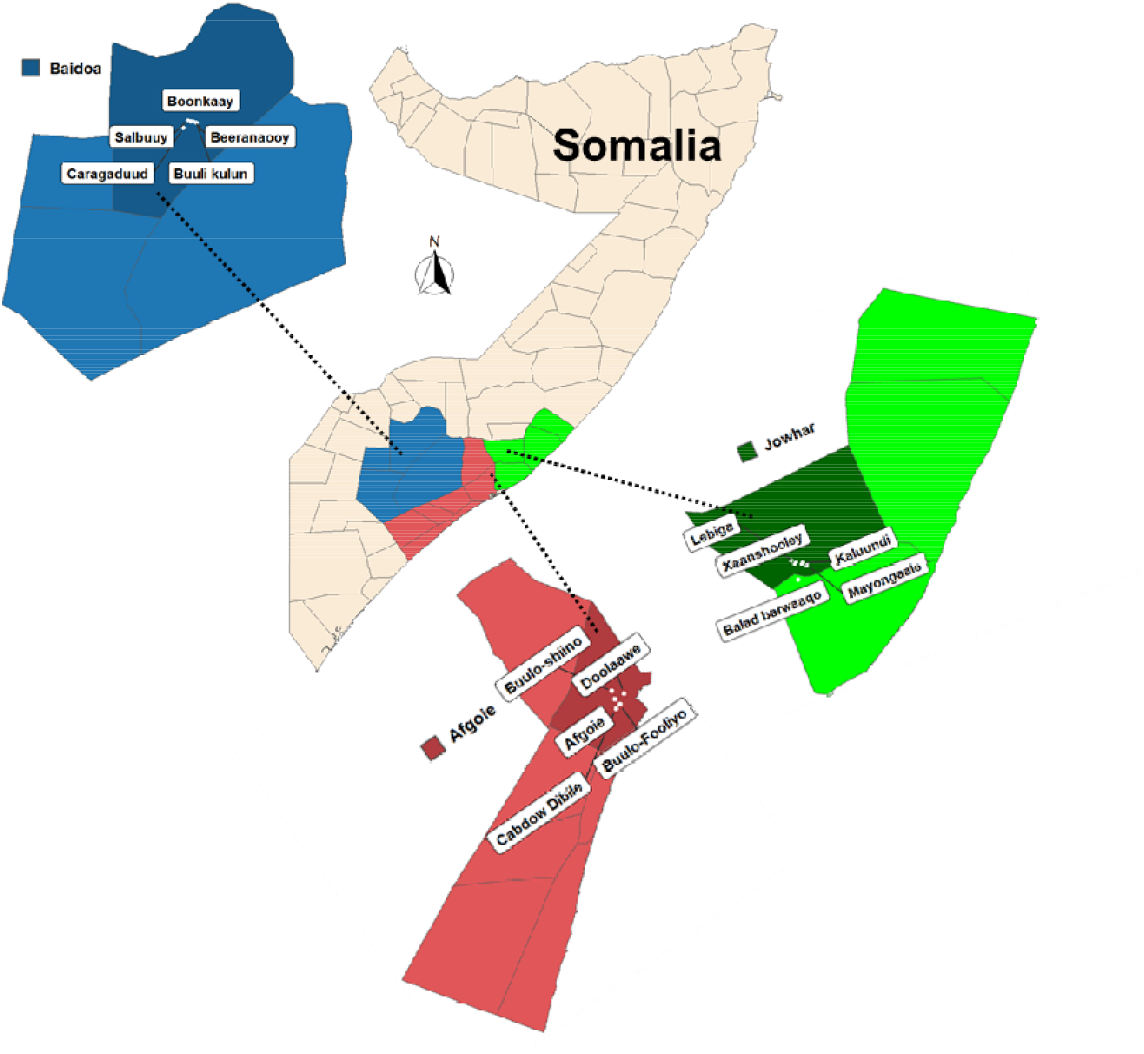
Map of the regions (provinces) in southern Somalia where farmer surveys were conducted. The three regions, Bay, Middle Shabelle, and Lower Shabelle, are shown on the main map in blue, green, and red, respectively. The dark-coloured segments in the map insets denote the selected districts within each region (Baidoa, Afgoie, and Jowhar). Villages where the farmer interviews took place are marked by white dots and labelled with their local names.

### Farmer surveys

A stratified random sampling approach was employed to ensure the representation of heterogeneous farming systems in the study regions. A total of ten farmers were randomly selected from each village, resulting in a sample size of 150 farmers (50 per district). Data were collected using a semi-structured electronic questionnaire designed and administered via the KoboToolbox platform (https://kf.kobotoolbox.org/). Trained enumerators conducted the interviews in Somali language. The questionnaire was organised into five sections: (a) farmer profile, (b) cowpea production system, (c) seed source and varietal awareness, (d) production constraints, and (e) farmer priorities for trait improvement. Initially, a pilot test was conducted with three farmers in each district to validate the questionnaire, after which a full survey was conducted from February to March 2025.

### Germplasm collection

To create a germplasm collection that captures Somalia’s cowpea genetic diversity, a nationwide cowpea seed collection was conducted from November 2024 to April 2025. A total of 60 seed samples were collected from major cities, small towns, remote villages, and farmers’ fields. The collection effort was primarily facilitated by volunteers recruited through social media platforms who dispatched seeds from various regions of the country. Upon receipt, the samples were stored in airtight containers and labelled with the collection site and a serial number. To ensure the genetic purity of the new collections, single seeds from each sample were grown in the field at Zamzam University of Science and Technology in Mogadishu.

In addition to the new collections from Somalia, pre-civil war cowpea collections were obtained from the International Institute for Tropical Agriculture (IITA), Austratian Grain Genebank (AGG), and USDA Germplasm Resources Information Network (USDA-GRIN). Furthermore, to broaden the genetic base of the core collection for breeding and genetic characterisation, additional lines from neighbouring countries, a subset of the IITA mini core (Fatokun et al., 2018), and lines with known desirable agronomic traits were included (see details in the Results section).

### Data analysis

The survey data were analysed using the R statistical environment. Descriptive statistics, including the percentage distributions of farmer responses, were computed and visualised using the base R functions and specialised R packages. On the other hand, to determine the main colour groups of the currently grown cultivars and those collected by Padulosi and Fiore (1989) just before the civil war in 1988-1989, seed coat colours were measured using the *Traitor* python package (Dayrell *et al*., 2023). For this analysis, at least 20 seeds were photographed for each accession, and RGB colour codes were obtained for each seed. The median RGB colour for each accession was then calculated and used for the PCA.

## RESULTS

### Cowpea production system in southern Somalia

To understand the current state of cowpea production in Somalia, farmer surveys were conducted across three major cowpea-producing regions: Bay, Middle Shabelle, and Lower Shabelle (Figure 1). These regions are representative of the common agroecological and livelihood systems in southern Somalia (Venema, 2007). A total of 150 farmers were interviewed, with nearly equal representation of males and females in Afgoie and Baidoa. However, the number of male respondents was relatively higher in Jowhar than in other regions (Figure 2A). Among the districts, farmers in Baidoa had, on average, twice as many years of growing cowpeas as their counterparts in Afgoie and Jowhar (Figure 2B). Similarly, the average farm size in Baidoa was larger (Figure 2C), despite the considerably lower cowpea yields reported by the farmers (Figure 2D). This is likely to compensate for the low productive per unit area. Likewise, due to the prevalence of agropastoralism in the Bay region (Longley et al., 2001a; World Bank/FAO, 2018), some farmers may require large fields to support fodder production during the dry season.

**Figure 2.**
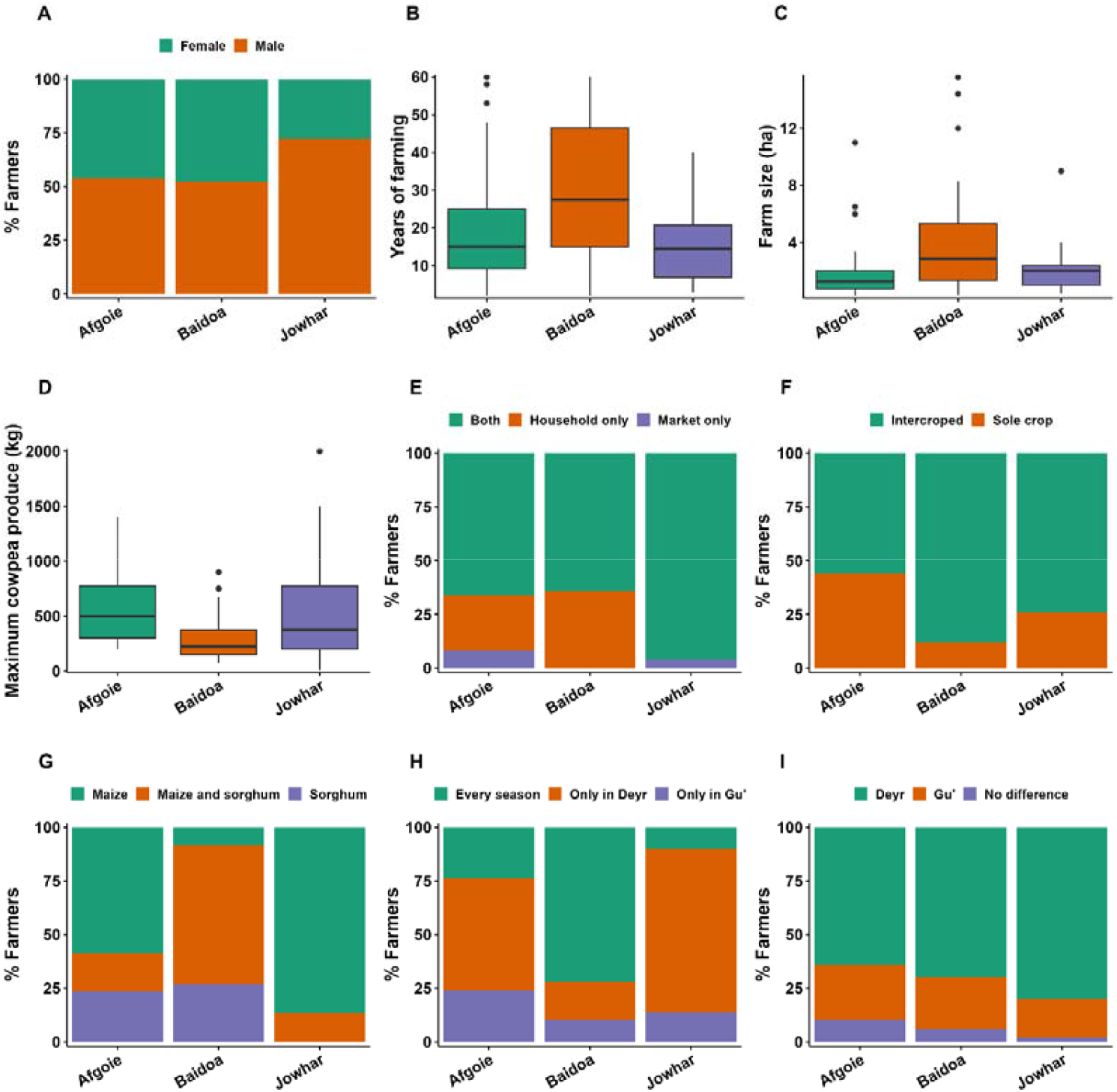
Farmer demographics and characteristics of cowpea production system in southern Somalia. Bar plots (A, E–I) represent the percentage of farmers responding to closed-ended questions on gender (A), main purpose of cowpea production (E), cultivation system (F), primary cereal intercrop (G), frequency of cultivation (H), and perceived optimal season for cowpea growth (I). Boxplots show years of experience in cowpea farming (B), farm size (C), and maximum cowpea yield during a good season (D).

Across the three districts, cowpea is primarily cultivated for both household consumption and commercial purposes. Notably, approximately 30% of farmers in Baidoa grow cowpea solely for subsistence (Figure 2E), highlighting its significance for household food security in this region. Intercropping was the predominant cowpea production system in all districts (Figure 2F). However, in the Afgoie district, approximately 45% of farmers reported cultivating cowpea as a sole crop. This trend is likely influenced by the district’s proximity to Mogadishu, the capital city, where there is a higher market demand for cowpea, thereby driving a shift towards sole cropping for commercial reasons. The three districts also differed in their cowpea-cereal combinations, with maize being the principal intercrop in Afgoie and Jowhar, while both maize and sorghum were used in Baidoa, likely planting sorghum during the shorter Deyr season and maize during the longer Gu’ season (Figure 2G). Furthermore, the difference between Baidoa and the other two districts was also evident in the main growing season of cowpea cultivation. In Baidoa, 75% of farmers reported growing cowpea in both the Deyr and Gu’ seasons, whereas in Afgoie and Jowhar, 50% and 70% of farmers, respectively, cultivated cowpea exclusively during the Deyr season (Figure 2H). The Deyr season (October–November) was regarded as the optimal period for cowpea production across all regions (Figure 2I). The preference for the Deyr season is likely because the end of the Gu season (around June) coincides with the onset of the *Hagaa* rains (Monsoon) along the Indian Ocean coast, which often complicate cowpea harvesting and promote the spread of diseases, thereby reducing productivity.

### Seed systems and varietal preferences

Farmers in the study area mainly obtained sowing seeds from either home-saved stocks or local grain markets (Figure 3A). In terms of locally grown cultivars, over 55% of farmers in Afgoie and Baidoa were not restricted to growing a particular variety, whereas more than 80% of farmers in Jowhar reported cultivating the same local variety each season (Figure 3B). This is despite the fact that at least three cowpea cultivars have been reported by farmers in each region (Figure 3C). In Baidoa, Longley et al. (2001a) reported three main cowpea varieties, namely *Bobodo*, a small-seeded and early maturing variety, *Degelo*, a large-seeded and late maturing, and *Abgalley*, which was introduced from Lower Shabelle region by NGOs. The role of NGOs in Somalia’s seed system was also evident in this study, with over 75% of the farmers in the three districts having received seeds from NGOs in the last five years (Figure 3D).

**Figure 3.**
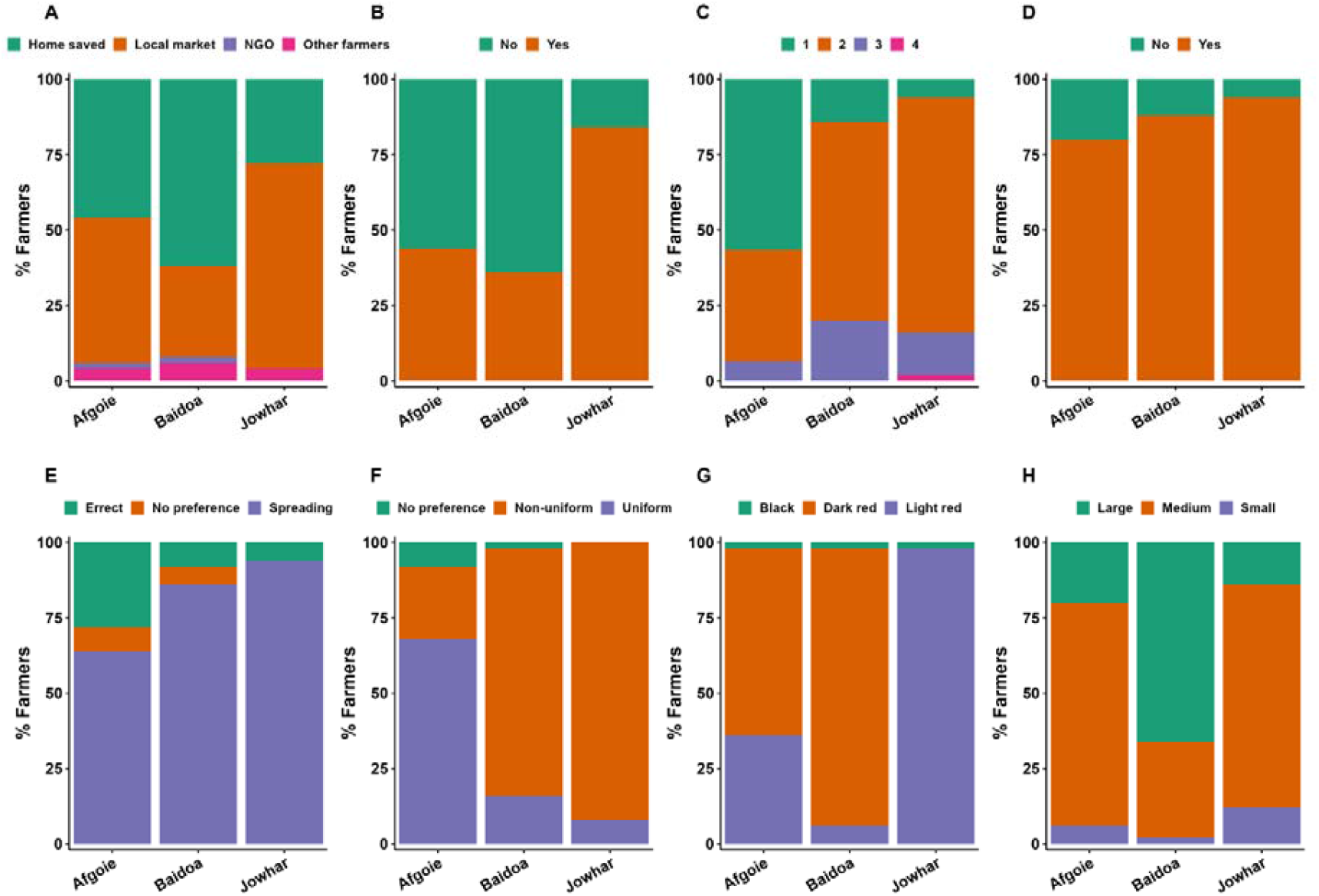
Sources of sowing seeds and varietal preferences among farmers in Baidoa, Afgoie, and Jowhar districts. The plots show sources of cowpea sowing seeds(A), consistency in cultivar selection (B), number of local cowpea cultivars known to farmers (C), receipt of sowing seeds from NGOs in the past five years (D), preferred cowpea growth habit (E), preferred maturity pattern (F), preferred seed colour (G), and size (H).

To establish the farmers’ preferences for cowpea varietal attributes, they were asked to choose between key agronomic traits related to plant architecture and seed attributes. In the three districts, farmers overwhelmingly favoured the spreading (prostrating) growth habit over the erect type (Figure 3E). Given that the spreading types are generally indeterminate and mature at different stages, farmers were further probed for their preference for uniformity in maturity. Unlike Jowhar and Baidoa, 65% of farmers in Afgoie preferred uniformly maturing cultivars (Figure 3F). This aligns with the relatively higher preference for the erect growth habit (Figure 3E) and the fact that 45% of farmers in Afgoie grow cowpea as a sole crop (Figure 2F). These findings underscore that cowpea production in the Afgoie district is more market-driven; thus, traits that facilitate commercial production, such as erect growth and uniform maturity, are preferred traits. Additionally, a region-specific preference for seed colour and size was identified (Figure 3G-H), with dark red being the desired colour in Afgoie and Baidoa, whereas light red varieties were favoured in Jowhar.

### Production constraints and priorities for genetic improvement

In Jowhar, Afgoie, and Baidoa, 86%, 62%, and 58% of the respondents, respectively, reported a declining trend in cowpea yields (Figure 4A). Furthermore, over 80% of farmers experienced complete crop failure at least once in the past (Figure 4B). However, the factors contributing to crop yield loss varied significantly among these regions. For example, drought was identified as the primary limiting factor in Baidoa, whereas diseases and pests were recognised as the main constraints by 56% and 42% of the farmers in Jowhar and Afgoie, respectively (Figure 4C). Although it was challenging to reconcile the various local names for the diseases and insect pests, the most frequently reported in these districts was aphids, referred to as *malabiso* in the local language, whereas leaf curl was the most common cowpea disease reported by farmers (data not shown).

**Figure 4.**
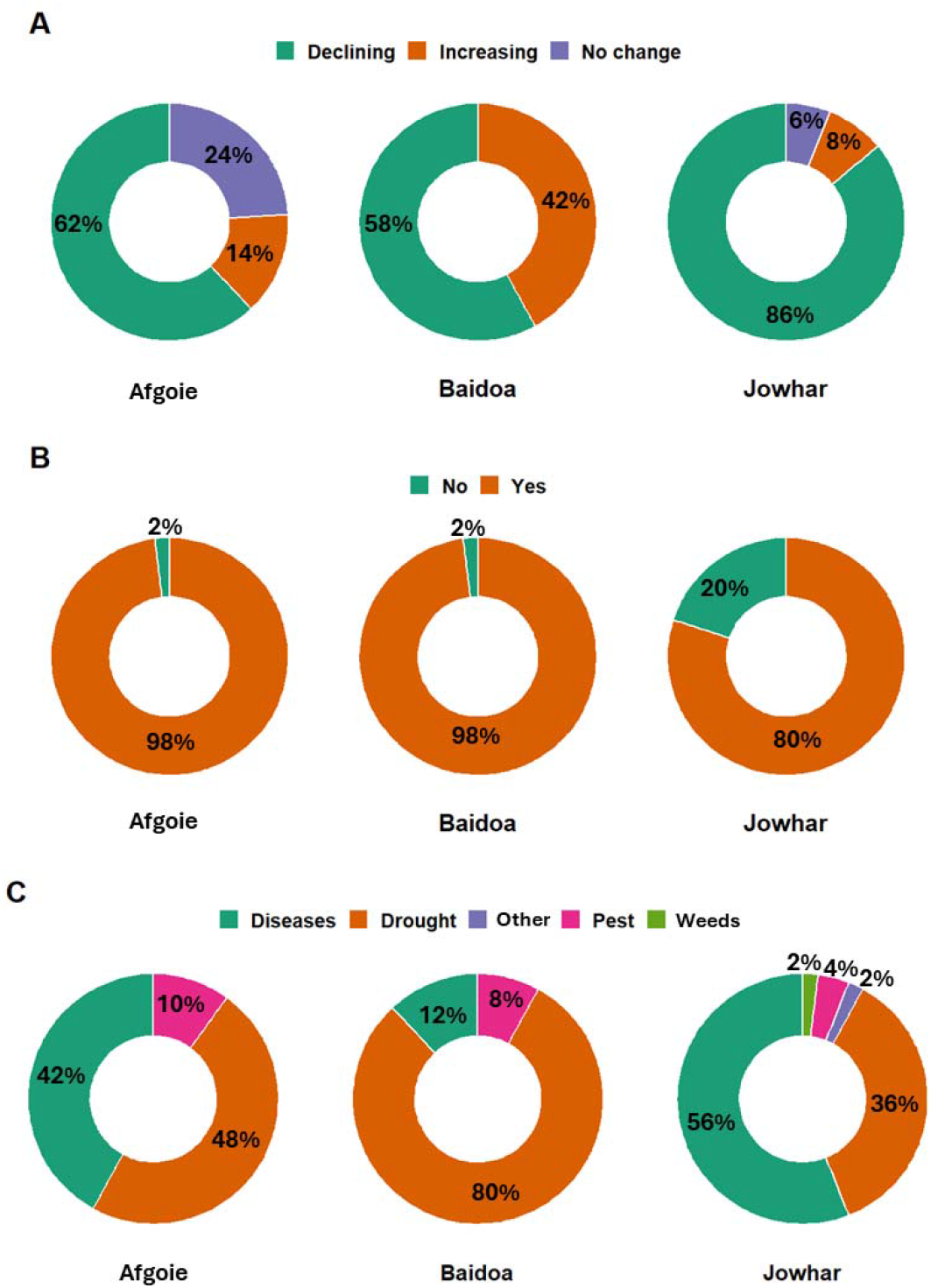
Farmers’ perspectives on the current trends in cowpea yields (A), their experiences with crop failure in recent years (B), and the primary factors affecting cowpea production (C).

To understand the priorities of farmers for cowpea genetic improvement, participants were asked to rank five agronomic and three quality traits in order of importance (Figure 5). Among the agronomic traits, early maturity was deemed highly significant in Baidoa, with 40% of the farmers identifying it as their foremost priority. This preference aligns with the region’s reliance on rain-fed crop production and the prevalence of drought as a primary production constraint. In this context, farmers appear to understand the importance of early flowering compared with drought tolerance as a trait, suggesting that early flowering to escape drought may be a more effective breeding target in this region. Conversely, in the Afgoie district, 55% of the farmers prioritised disease and pest resistance. This finding is consistent with the 52% that reported diseases and pests as the main constraints to cowpea production in Afgoie (Figure 5C). Regarding quality traits, reducing digestive discomfort, which is caused by higher concentrations of raffinose family oligosaccharides, and fast cooking time were considered of higher importance by farmers in Jowhar and Baido, respectively.

**Figure 5.**
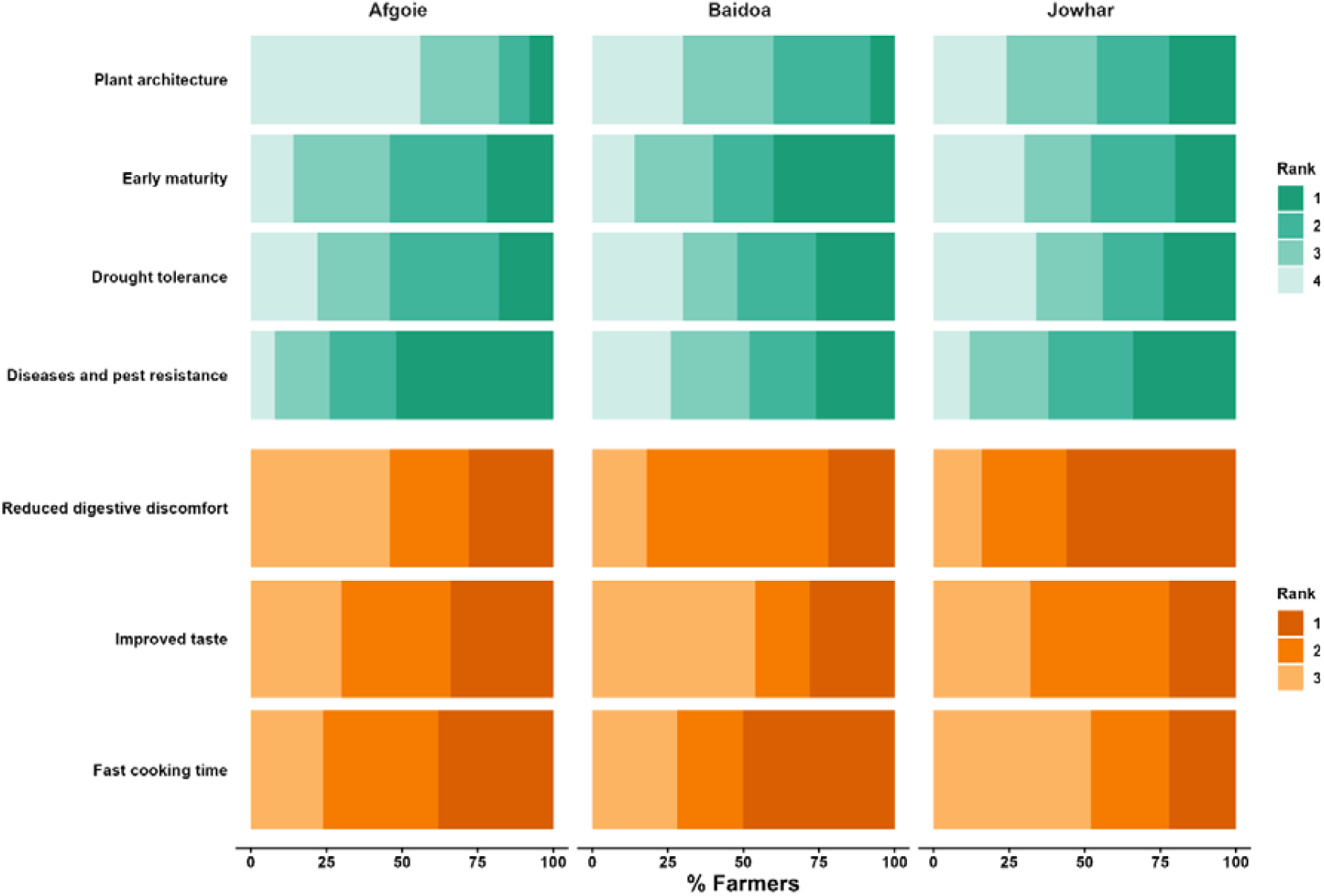
Agronomic (top panel) and quality traits (bottom panel) of cowpea and their farmer-prioritised ranking for improvement in three regions of southern Somalia. The shades represent the priority levels, with darker colours indicating a higher priority for improvement.

### Creating core germplasm collection

Somalia’s agroecology is highly heterogeneous and encompasses arid and semi-arid rangelands, fertile river valleys, and extensive coastal zones, which in turn shape distinct livelihood systems, including pure pastoralism, agro-pastoralism, and rain-fed and irrigated farming systems (Figure 6). Therefore, the cowpea germplasm collections by Padulosi and Fiore (1989) and ours in 2024–2025, span a wide range of ecosystems, extending from the agro-pastoral areas of the northwest (Somaliland) through the central ‘cowpea belt’ to the agro-pastoral and riverine lands of southern Somalia (Figure 6). Furthermore, the two collections do not overlap spatially and temporally, the two collections together can be considered to represent the core genetic diversity of cowpea in Somalia. Additionally, among the historical cowpea accessions, five were obtained from the Australian Grain Genebank, which dates back to 1973, and a single accession from USDA-ARS-GRIN, with an acquisition date of 1958 (Table 1). To further broaden the genetic base and facilitate fast-tracking cowpea breeding, we included 103 accessions from different parts of the world, including neighbouring countries and West African countries with arid climates, including Mali, Niger and Benin. Furthermore, the collection included 21 accessions with reported resistance to biotic and abiotic stresses (USDA-ARS Plant Genetic Resources Conservation Unit, 2020). The core collection comprised 250 accessions.

**Table 1.** Types, sources, and number of accessions constituting the new core germplasm for cowpea genetic improvement in Somalia.

| Collections | Source | Number of accessions | Year of collection |
| --- | --- | --- | --- |
| **1. Somali cowpea collections** |  |  |  |
| New collection | Somalia | 60 | 2024-2025 |
| Historical collection (1) | IITA | 50 | 1988-1989 |
| Historical collection (2) | AGG | 5 | 1973 |
| Historical collection (3) | USDA-GRIN | 1 | 1958 |
| **2. Accessions from neighbouring countries** |  |  |  |
| Kenya | IITA | 5 | NA |
| Ethiopia | IITA | 5 | NA |
| **3. Accessions with known desirable traits** |  |  |  |
| Drought tolerant | IITA & USDA-GRIN | 7 | NA |
| Disease and pest resistant | IITA | 14 | NA |
| **4. A subset of Global cowpea diversity** |  |  |  |
| IITA mini core | IITA | 103 | NA |
| **Total** |  | **250** |  |

IITA=International Institute for Tropical Agriculture; AGG=Australian Grain Genebank; USDA=United States Department of Agriculture

**Figure 6.**
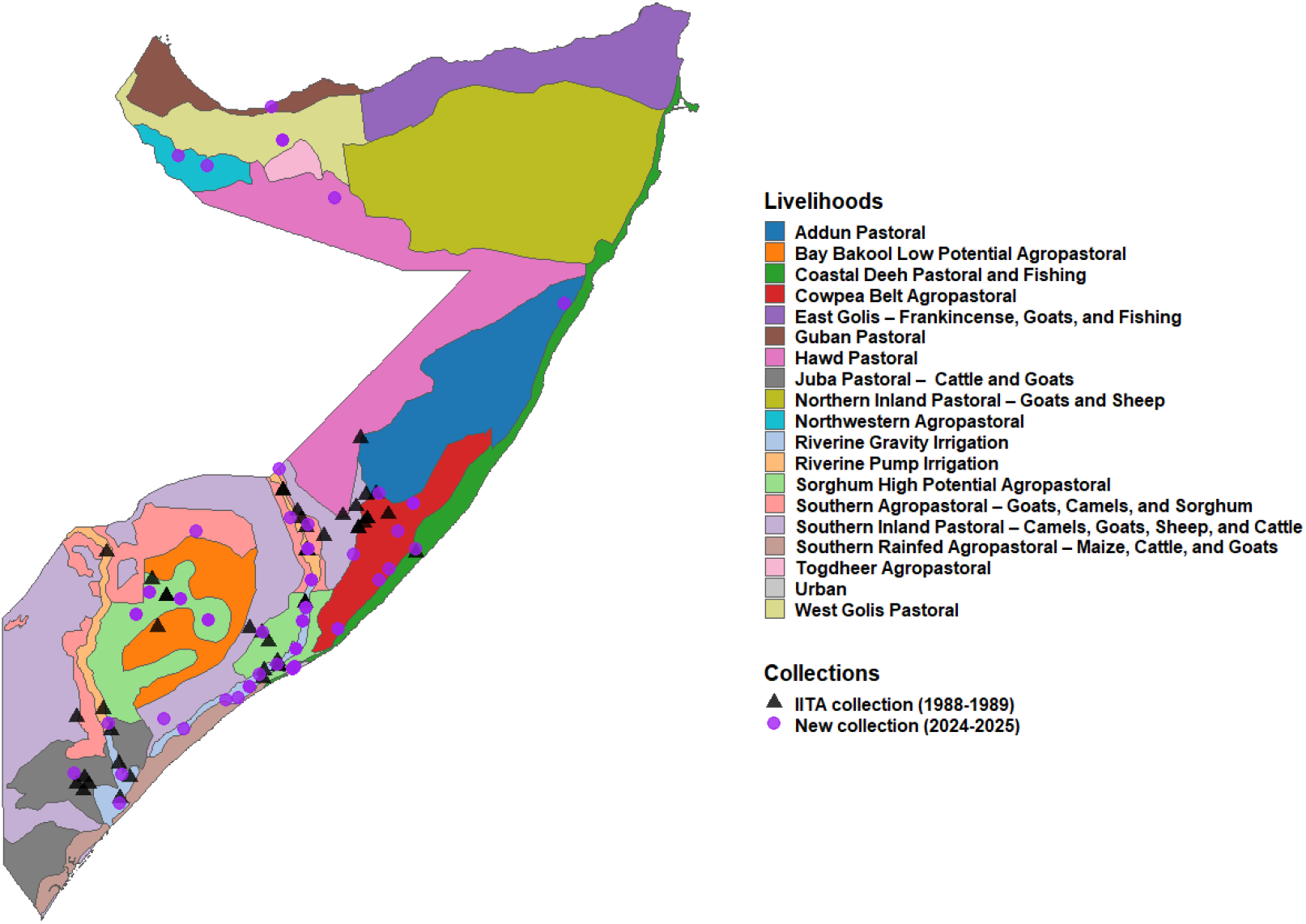
Core cowpea germplasm collection from regions spanning different livelihood systems in Somalia. Black triangles represent IITA collections conducted between 1988 and 1989, and purple circles indicate locations of the new collections from 2024 to 2025. The map of livelihood zones was drawn based on data sourced from FEWS NET (https://fews.net).

Seed coats are considered the most important variety descriptor and determine regional preferences for cowpea varieties *(Boukar et al., 2019*; *Murdock et al., 2013)*. Therefore, to obtain information on the main colours of cowpea cultivars in Somalia, accessions were grouped based on visual and quantitative measurements of seed colour intensity (Figure 7). Overall, red colour, ranging from light to dark, was the dominant cowpea colour in both the old and new collections. Eleven and nine accessions in the old (Figure 7A) and new (Figure 7B) collections had creamy colours, of which nearly all were from the central to northern parts of Somalia, including El-bur, Bargan, Galhareri, and El-dher in the Galgudud region. The distribution of seed coat colour aligns closely with local culinary practices and consumer preferences for cowpea in Somalia. In major cities of southern Somalia, cowpea is typically cooked together with rice or maize, and red-seeded cultivars are preferred because they give the characteristic red colour to the popular dinner dish known as *Anbuulo*. However, in the central regions, where cowpea is consumed alone as boiled or *Falfaliir*, a dish prepared from roasted, dehulled, and then boiled cowpea, the light creamy colours are desired. Finally, there were few accessions with notable colours including black-eyed accession (Som-Vu054) from Gal Hareeri, black seeded TVu-16069 from Afgoie, and large-seeded accession (Som-Vu051) from Awdheegle.

**Figure 7.**
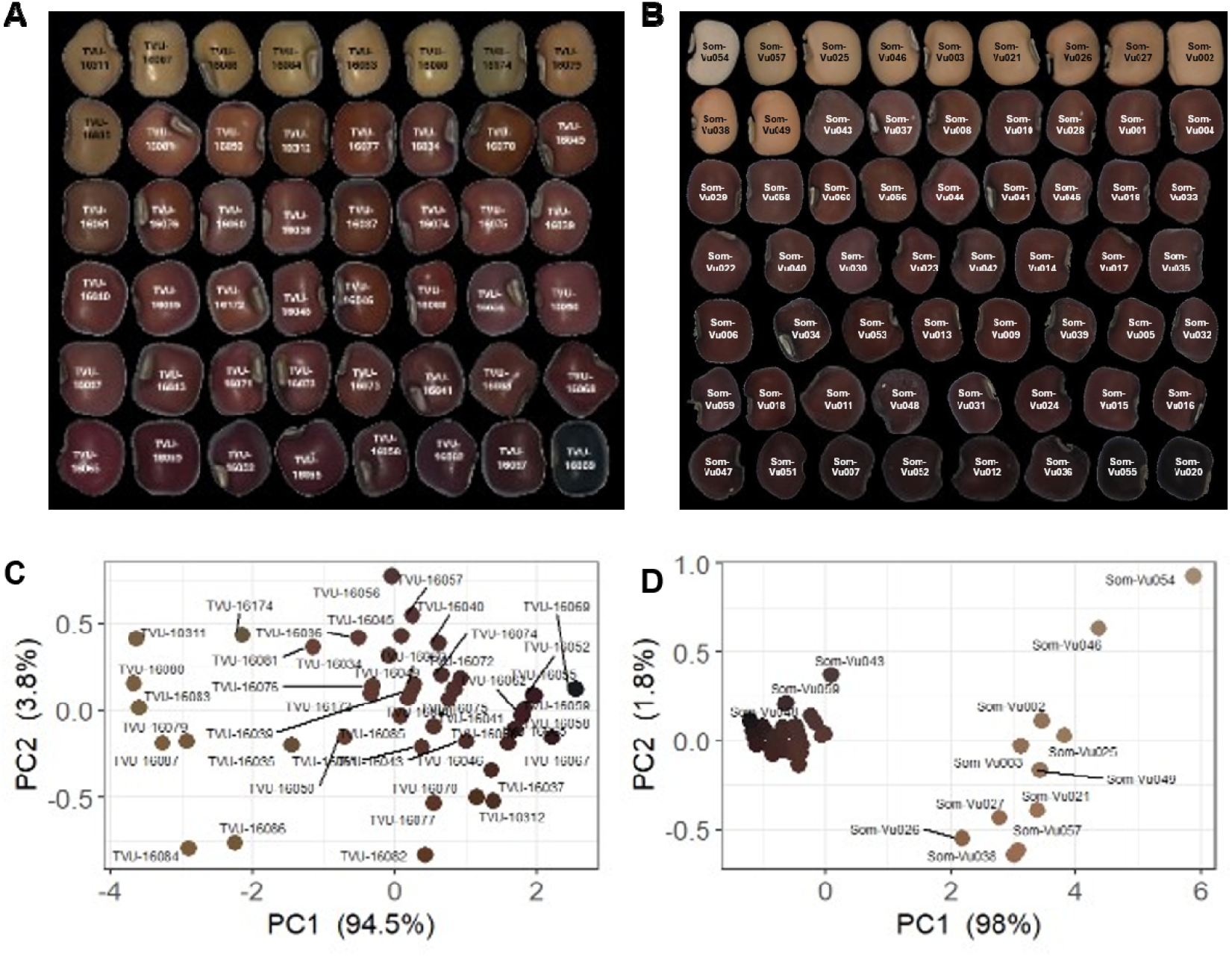
Seed coat colour variation in Somalia’s historical and currently grown cowpea cultivars. Each seed image in A and B represents the seed colour of individual accessions in the IITA collection (1988-1989) and the new collections of this study (2024-2025), respectively. The Principal Component Analysis (PCA) plots are based on the median RGB colour of at least 20 seeds of each accession, and represent the collections of IITA (C) and this study (D)

It is worth noting that the data on seed colour primarily reflects the preferred colour of cowpea and does not adequately capture the actual diversity in the population. For example, within the predominant red colour group in the currently cultivated varieties, there is notable phenotypic variation in other traits, including plant architecture, ranging from extremely spreading to semi-erect, as well as differences in earliness, stem pigmentation intensity, and susceptibility to diseases, particularly leaf curl mosaic virus (Figure 8). Consequently, a more comprehensive genetic analysis is necessary to elucidate the overall population structure and extent of genetic diversity loss or gain in the post-conflict cowpea population in Somalia.

**Figure 8.**
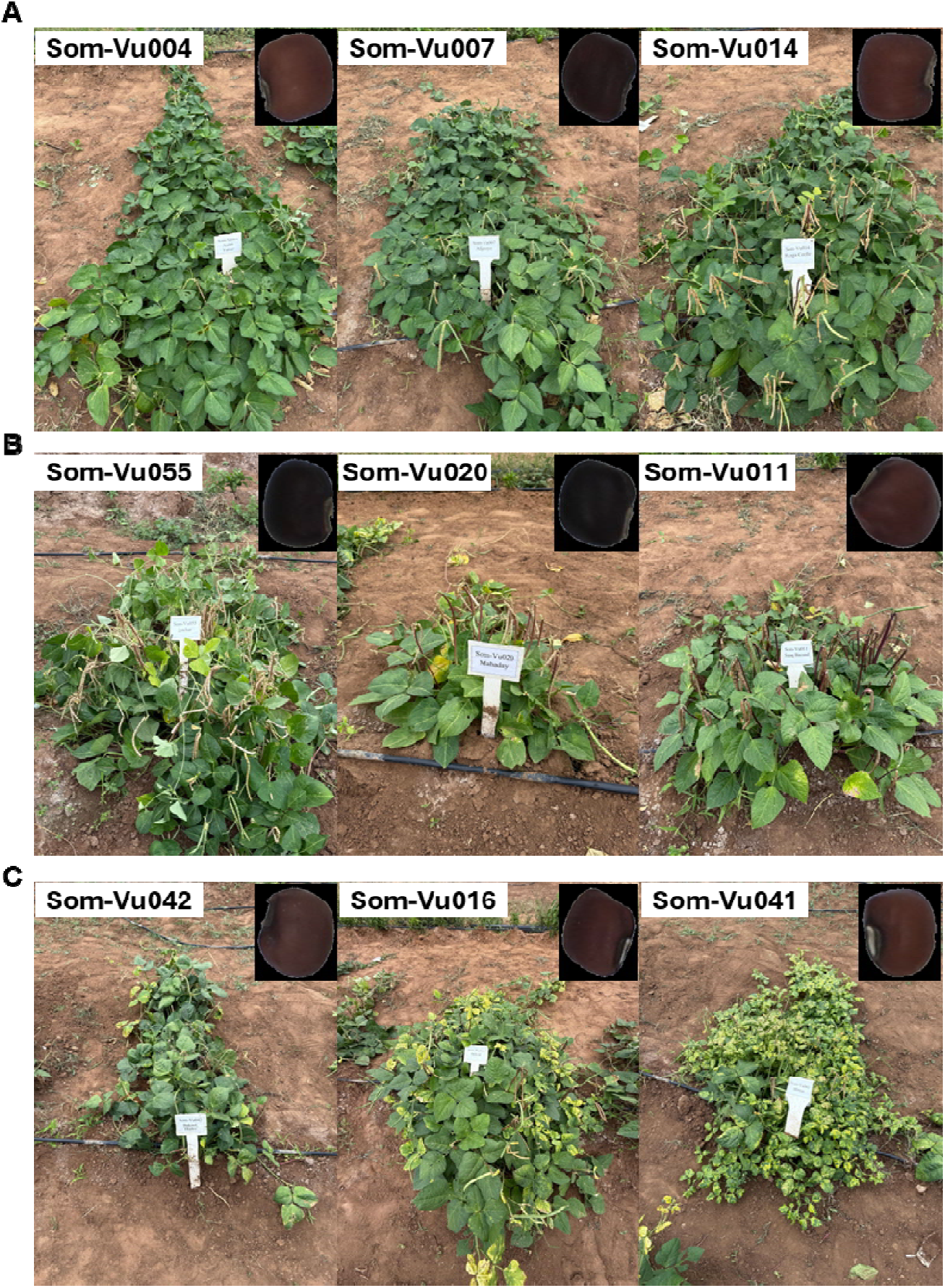
Phenotypic variation within red-coloured cowpea cultivars currently grown in Somalia. A. represents highly spreading cultivars with observable differences in their pod-setting and maturity. The cultivars in B show different degrees of stem erectness and pigmentation, and C are accessions infected by leaf curl mosaic virus.

## DISCUSSION

Apart from a few reports by international development organisations (Longley et al., 2001a; World Bank/FAO, 2018), there are no systematically collected data on the status of Somalia’s major food crops after more than three decades of political instability and recurrent conflict. Therefore, this study represents the first comprehensive report on cowpea production, the main constraints, and genetic resources in the country.

The three regions selected for farmer surveys were broadly representative of south-central Somalia, where cowpea is mainly produced, ensuring that the findings are relevant to the majority of cowpea production zones. Furthermore, while the sample size of 150 farmers was relatively modest compared to similar studies, such as 260 respondents in Jinbaani et al. (2023) and 320 in Martey et al. (2022), the study provides a robust description of the current cowpea production system in Somalia and identifies clear priorities for agronomic and quality improvements.

The study revealed regional differences in cowpea production systems, cultivar preferences, major production constraints, and traits of priority for breeding. For example, in Baidoa, indeterminate, spreading and red-seeded cultivars are grown for intercropping with sorghum. In such a system, an erect, black-seeded variety, even if agronomically superior, is unlikely to be adopted because of its incompatibility with established production systems and market preferences. On the other hand, in Afgoie district, our findings suggest that new cowpea varieties with desirable traits, including erect growth habit, uniform maturity, and other quality traits, could facilitate viable commercial cowpea production to meet the market demands in the capital city, Mogadishu. Such geographically defined preferences have been reported among cowpea farmers in other parts of Africa (Ishikawa et al., 2020; Jinbaani et al., 2023; Martey et al., 2022), indicating the need for breeding programs and seed interventions tailored to specific agroecological and socioeconomic conditions. The historical case reported by Rees et al. (1991), in which the early maturing semi-erect variety from IITA was not preferred by farmers due to its undesirable seed colour, reinforces the importance of combining agronomic performance with farmer and market preferences.

To meet current and future cowpea breeding priorities in Somalia, we assembled a core germplasm collection with a broad genetic base, encompassing local, regional, and global collections. It is worth noting that East Africa is recognised as one of the centres of origin of cowpea (Herniter et al., 2020; Xiong et al., 2016), and the arid and semi-arid zones of the Horn of Africa may harbour unique genetic variants that are valuable for genetic improvement of drought tolerance, pest and disease resistance, and desirable grain qualities. However, decades of civil strive, which have disproportionately affected farming communities (Majid & McDowell, 2012; Maxwell & Fitzpatrick, 2012), may have significantly eroded some of the local crop diversity. For instance, during the Rwandan civil war, which lasted for a few months, one-third of farmers reported losing their local bean cultivars (Sperling, 2001). Other factors, such as drought and pest damage, can also drive crop variety losses (Horn & Shimelis, 2020). Therefore, it is plausible that Somalia’s cowpea gene pool has been narrowed not only because of conflict and farmer displacement but also because of climatic shocks and the introduction of external seeds through aid programs.

The cowpea genetic resources established in this study can be exploited in various ways to address the urgent production constraints and market demands in Somalia. Core collection can serve as foundational material for pre-breeding and cultivar development programs. This can be achieved by field screening for yield potential, drought tolerance, and resistance to insect pests and fungal, bacterial, and viral diseases. Furthermore, it will be crucial to assess the diversity for attributes that directly influence consumer acceptance and marketability, including sensory, nutritional, and culinary qualities. Additionally, given Somalia’s agroecological heterogeneity and region-specific varietal preferences, participatory varietal selection (PVS) approaches will be critical to ensure that newly developed cultivars align with both agronomic requirements and sociocultural preferences. Finally, this collection is a valuable research material for training the next generation of crop scientists and plant breeders in Somalia.

## CONCLUSION

This study is the first report on the state of cowpea production in Somalia, and provides insights into the production practices, main production constraints, and farmers’ perspectives on the traits of high priority for genetic improvement. Furthermore, a comprehensive collection of cowpea germplasm was initiated. Overall, this study establishes a foundation for targeted genetic improvement of cowpea in Somalia and outlines a pathway for developing climate-smart cultivars that combine desirable agronomic and sensory qualities. Such varieties have the potential to enhance the food security and livelihoods of smallholder farmers, while providing sustainable, affordable sources of plant-based protein to meet the dietary needs of the country’s rapidly growing urban population.

## Author Contributions

Ahmed O. Warsame conceptualised the research project, designed the survey, analysed and drafted the manuscript. Yahye A. Isse and Abdirizak Mohamed Sh. Abdi supervised and conducted the surveys, and conducted and coordinated the local germplasm collection. All authors contributed to the writing, review, and revision of the final manuscript.

## Acknowledgments

We acknowledge the many people who contributed to the collection of cowpea germplasm, including Ali Omar Warsame, who played a crucial role in facilitating sample shipments from remote areas in central Somalia. We would also like to thank the staff at the Ministry of Agriculture and Irrigation of the Federal Republic of Somalia for their contribution to the sample collection efforts. We also acknowledge the staff members of the Zamzam University of Science and Technology in Jowhar and Baidoa for their contribution to the farmer surveys. This research was supported by the Allan and Gill Gray Foundation Grant to the Cambridge University Crop Science Centre (grant no G118688).

